# Pan-cancer image-based detection of clinically actionable genetic alterations

**DOI:** 10.1101/833756

**Authors:** Jakob Nikolas Kather, Lara R. Heij, Heike I. Grabsch, Loes F. S. Kooreman, Chiara Loeffler, Amelie Echle, Jeremias Krause, Hannah Sophie Muti, Jan M. Niehues, Kai A. J. Sommer, Peter Bankhead, Jefree J. Schulte, Nicole A. Cipriani, Nadina Ortiz-Brüchle, Akash Patnaik, Andrew Srisuwananukorn, Hermann Brenner, Michael Hoffmeister, Piet A. van den Brandt, Dirk Jäger, Christian Trautwein, Alexander T. Pearson, Tom Luedde

**Affiliations:** Department of Medicine III, University Hospital RWTH Aachen, Aachen, Germany; German Cancer Consortium (DKTK), Heidelberg, Germany; Applied Tumor Immunity, German Cancer Research Center (DKFZ), Heidelberg, Germany; Department of Surgery and Transplantation, University Hospital RWTH Aachen, Aachen, Germany; Department of Surgery, NUTRIM, School of Nutrition and Translational Research in Metabolism, Maastricht University, Maastricht, The Netherlands; Institute of Pathology, University Hospital RWTH Aachen, Aachen, Germany; Department of Pathology, GROW School for Oncology and Developmental Biology, Maastricht University Medical Center+, Maastricht, The Netherlands; Pathology & Data Analytics, Leeds Institute of Medical Research at St James’s, University of Leeds, Leeds, UK; MRC Institute of Genetics and Molecular Medicine, University of Edinburgh, Edinburgh, UK; Department of Pathology, University of Chicago Medicine, Chicago, IL, USA; Department of Medicine, University of Chicago Medicine, Chicago, IL, USA; Department of Medicine, University of Illinois – Chicago, Chicago, IL, USA; Division of Clinical Epidemiology and Aging Research, German Cancer Research Center (DKFZ), Heidelberg, Germany; Division of Preventive Oncology, German Cancer Research Center (DKFZ) and National Center for Tumor Diseases (NCT), Heidelberg, Germany; Department of Epidemiology, GROW School for Oncology and Developmental Biology, Maastricht University Medical Center+, Maastricht, The Netherlands; Division of Gastroenterology, Hepatology and GI Oncology, University Hospital RWTH Aachen, Aachen, Germany

## Abstract

Precision treatment of cancer relies on genetic alterations which are diagnosed by molecular biology assays.^1^ These tests can be a bottleneck in oncology workflows because of high turnaround time, tissue usage and costs.^2^ Here, we show that deep learning can predict point mutations, molecular tumor subtypes and immune-related gene expression signatures^3,4^ directly from routine histological images of tumor tissue. We developed and systematically optimized a one-stop-shop workflow and applied it to more than 4000 patients with breast^5^, colon and rectal^6^, head and neck^7^, lung^8,9^, pancreatic^10^, prostate^11^ cancer, melanoma^12^ and gastric^13^ cancer. Together, our findings show that a single deep learning algorithm can predict clinically actionable alterations from routine histology data. Our method can be implemented on mobile hardware^14^, potentially enabling point-of-care diagnostics for personalized cancer treatment in individual patients.

---

Clinical guidelines recommend molecular testing of tumor tissue for most patients with advanced solid tumors. However, in most tumor types, routine testing includes only a handful of alterations, such as KRAS, NRAS, BRAF mutations and microsatellite instability (MSI) in colorectal cancer. While new studies identify more and more molecular features of potential clinical relevance, current diagnostic workflows are not designed to incorporate an exponentially rising load of tests. For example, in colorectal cancer, previous studies have identified consensus molecular subtypes (CMS) as a candidate biomarker, but sequencing costs preclude widespread testing.

While comprehensive molecular and genetic tests are hard to implement at scale, histological images stained with hematoxylin and eosin (H&E) are ubiquitously available. We hypothesized that these routine images contain information about established and candidate biomarkers and thus could be used for rapid pre-screening of patients, potentially alleviating the load of molecular assays. To test this, we developed, optimized and validated a deep learning algorithm to determine molecular features directly from histology images. Deep learning with convolutional neural networks has been used for tissue segmentation in cancer histology^15–17^ or detecting molecular changes in circumscribed use cases in a single tumor type^18–22^, but our aim was to use deep learning in a pan-molecular pan-cancer approach. Our method is a ‘one-stop-shop’ workflow: we collected large patient cohorts for individual tumor types, partitioning each cohort into three groups for cross-validation (Fig. 1a). Whole slide images were tessellated into an image library of smaller tiles^20,21^ which were used for deep transfer learning (Fig. 1b). We chose prediction of microsatellite instability (MSI) in colorectal cancer as a clinically relevant benchmark task^20^ and sampled a large hyperparameter space with different commonly used deep learning models^16,18,20,21^. Unexpectedly, ‘inception’^23^ and ‘resnet’^24^ networks, which had been the previous defacto standard, were markedly outperformed by ‘densenet’^25^ and ‘shufflenet’^14^ architectures, the latter demonstrating high accuracy at a low training time (raw data in Suppl. Table 1, N=426 patients in the “Cancer Genome Atlas” [TCGA] cohort). Shufflenet is optimized for mobile devices, making this deep neural network architecture attractive for decentralized point-of-care image analyses or direct implementation in microscopes^26^. We trained a shufflenet on N=426 patients in the TCGA-CRC cohort^20^ and validated it on N=379 patients in the DACHS cohort^20^ cohort, reaching an AUC of 0.89 [0.88; 0.92] (Fig. 1d). This represents a marked improvement over the previous best performance of 0.84 in that dataset^20^. Subsequently, we tested the full workflow in breast cancer for detection of standard molecular pathology features which are usually measured by immunohistochemistry: Estrogen [ER] and progesterone [PR] receptor status and HER2 status were highly significantly detectable from histology alone, reaching AUCs of up to 0.82 in a three-fold patient-level cross-validation (Fig. 1e).

**Fig. 1:**
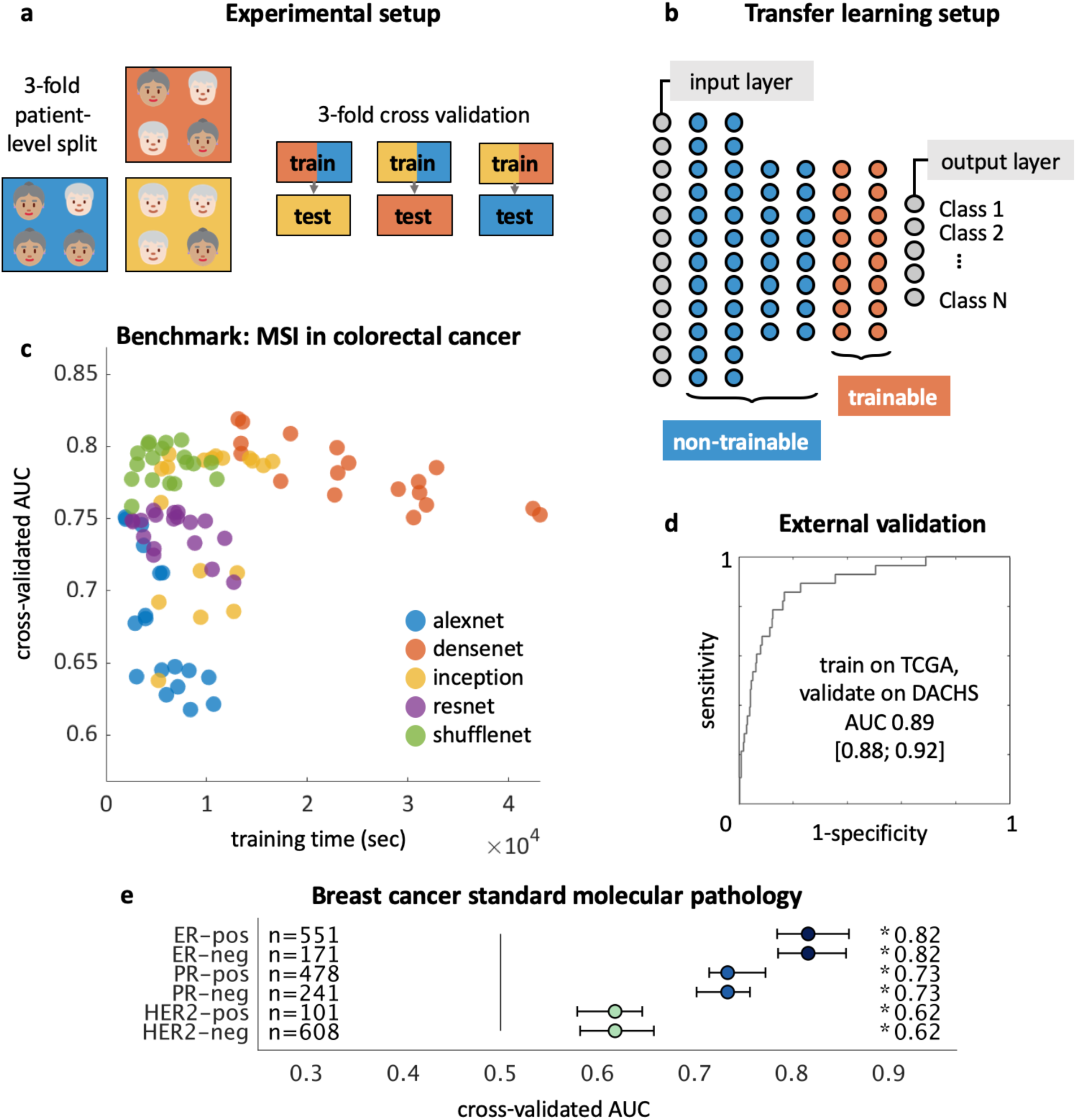
Transfer learning workflow for histology images. (**a**) Patient cohorts are split into three partitions for cross-validation of deep classifiers (**b**) Pre-trained networks re re-trained with only the deepest layers trainable, speeding up computation while enabling state-of-the-art performance. (**c**) A hyperparameter sweep with multiple networks shows that shufflenet consistently yields high accuracy and speed for detection of microsatellite instability (MSI) in colorectal cancer (N=426 patients), raw data in Suppl. Table 1. (**d**) External validation of the best shufflenet on the DACHS cohort (N=379 patients). (**e**) Validation of the workflow by prediction of estrogen receptor (ER), progesterone receptor (PR), HER2 status and tumor mutational burden (TMB) in breast cancer, assessed by cross-validated area under the receiver operating curve (AUC).

Having optimized our method in these use cases, we applied it to more than 4000 patients across ten of the most prevalent solid tumor types from the TCGA reference database. We aimed to predict all clinically and/or biologically relevant mutations with a prevalence above 2% and affecting at least four patients. The list of candidate mutations (Suppl. Table 2) also included all point mutations targetable by FDA-approved drugs (www.oncokb.org). We found that in multiple major cancer types, the genotype of point mutations was predictable directly from images. For example, in lung adenocarcinoma (TCGA-LUAD^8^, N=464 patients), significant AUCs were achieved for TP53 mutational status (AUC 0.71, Fig. 2a) and EGFR mutational status (AUC 0.60), which is targetable by clinically approved treatments. Also in colon and rectal cancer (TCGA-COAD and TCGA-READ^27^, N=590 patients), standard-of-care genetic biomarkers^28^ BRAF (AUC 0.66) and KRAS (AUC 0.60) were significantly detectable, as were oncogenic driver mutations linked to tumor aggressiveness, including CDC27^29^ (AUC 0.70, Fig. 2b). Similarly, in breast cancer (TCGA-BRCA^5^, N=1007 patients), gene mutations of TP53 (AUC 0.75), MAP2K4 (which is a potential biomarker for response to MEK inhibitors^30^, AUC 0.66) as well as PIK3CA (which is directly targetable by a small molecule inhibitor^31^, AUC 0.63) were significantly detectable (Fig. 2c). In gastric cancer (TCGA-STAD^13^, N=363 patients), mutations of MTOR – a candidate for targeted treatment^32^ – were significantly detectable with a high AUC of 0.80 (Fig. 2d) as were a range of driver mutations including BRCA2 (AUC 0.67), PTEN (AUC 0.66), PIK3CA (AUC 0.65) among others. In head and neck squamous cell carcinoma (TCGA-HNSC^7^, N=424 patients), genotype of CASP8, which is linked to resistance to cell death^33^, was significantly detected with a high AUC of 0.72 (Suppl. Fig. 1a). In other tumor types such as melanoma (TCGA-SKCM^12^, N=429 patients), or lung squamous cell carcinoma (TCGA-LUSC^9^, N=412 patients), few mutations were significantly detected (Suppl. Fig. 1b-c). Lung squamous cell carcinoma is known for its difficulty in molecular diagnosis and few molecularly or genetically targeted treatment options even in clinical trials. Thus, it is plausible that tumor histomorphology was not well correlated to mutations. In pancreatic adenocarcinoma (TCGA-PAAD^10^, N=166 patients), identifying KRAS wild type patients is of high clinical relevance because these patients are potential candidates for targeted treatment. Our method significantly identified KRAS genotype with AUC 0.66 (Suppl. Fig. 1d). Lastly, in prostate cancer (TCGA-PRAD^11^, N=402 patients), our method detected targetable mutations from histology – most remarkably PIK3CA, which was significantly detected with an AUC of 0.75 (Suppl. Fig. 1e). Furthermore, CDK12, which is linked to immune evasion in prostate cancer^34^ was detected with an AUC of 0.71. Together, these data show that deep learning can detect a wide range of targetable and potentially targetable point mutations directly from histology across multiple prevalent tumor types.

**Fig. 2:**
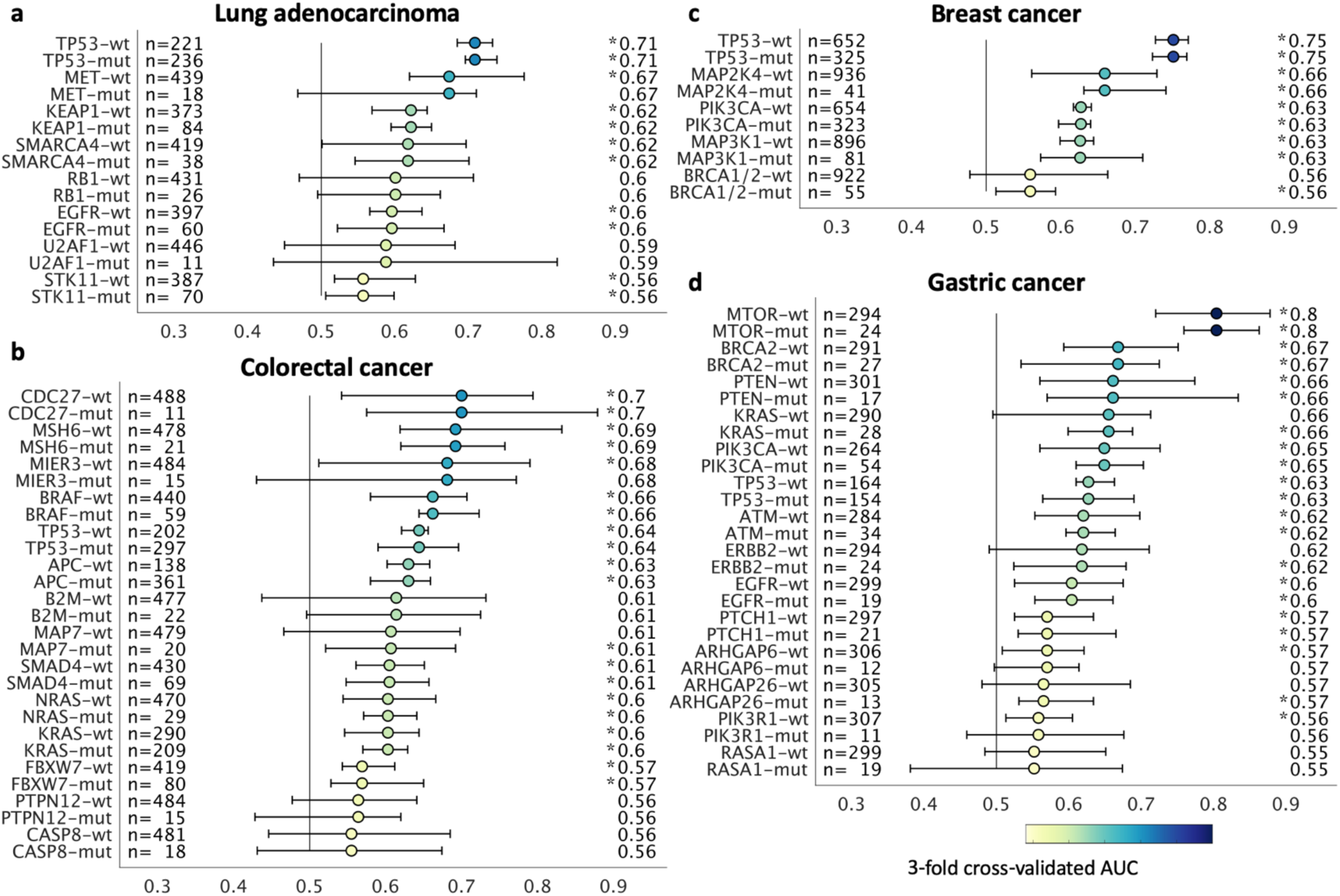
Prediction of point mutations directly from histological images. Deep networks predicted genotype directly from histological images in (**a**) lung adenocarcinoma, (**b**) colorectal, (**c**) breast cancer and (**d**) gastric cancer. Patient cohorts were randomly split for cross validation and classifiers were assessed by the area under the receiver operating curve (AUC, horizontal axis) with a 95% bootstrapped confidence interval. Genotype was predicted from histology with a high AUC for multiple clinically actionable mutations. (*) denotes all cases where the lower confidence bound exceeds a random classifier (AUC 0.5). “n” denotes the number of patients. Mutations with an AUC<0.55 are not shown. For a full list of all tested alterations, see Suppl. Table 2.

Next, we applied our method to a broader set of molecular signatures beyond single mutations. We chose features with known biological and potential clinical significance which are currently not part of clinical guidelines in most solid tumors. A major group of such features are immune-related gene expression signatures^3^ of CD8-positive lymphocytes, macrophages, proliferation, interferon-gamma (IFNg) signaling and transforming growth factor beta (TGFb) signaling. These biological processes are involved in response to cancer treatment, including immunotherapy. Detecting their morphological correlates in histology images could facilitate the development of more nuanced treatment strategies. Indeed, in lung adenocarcinoma signatures of proliferation, macrophage infiltration and T-lymphocyte infiltration were significantly detectable from images with high AUCs (Fig. 3a). Similarly, significant AUCs for these biomarkers were achieved in colorectal cancer (Fig. 3b), breast cancer (Fig. 3d) and gastric cancer (Fig. 3d). In gastric cancer, we additionally investigated a signature of stem cell properties (stemness) which was highly detectable in images (AUC 0.76, Fig. 3d). Recent studies have clustered tumors into comprehensive ‘immune subtypes’^3^, but again this classification system relies on deep molecular profiling unavailable in a clinical setting. We found that our method could detect these immune subtypes with up to AUC 0.75 in lung adenocarcinoma (Fig. 3a), up to AUC 0.72 in colorectal cancer (Fig. 3b) and up to AUC 0.71 in breast cancer (Fig. 3c). Together, these findings show that immunological processes that are quantifiable by molecular profiling are also accessible to deep-learning-based histology image analysis.

**Fig. 3:**
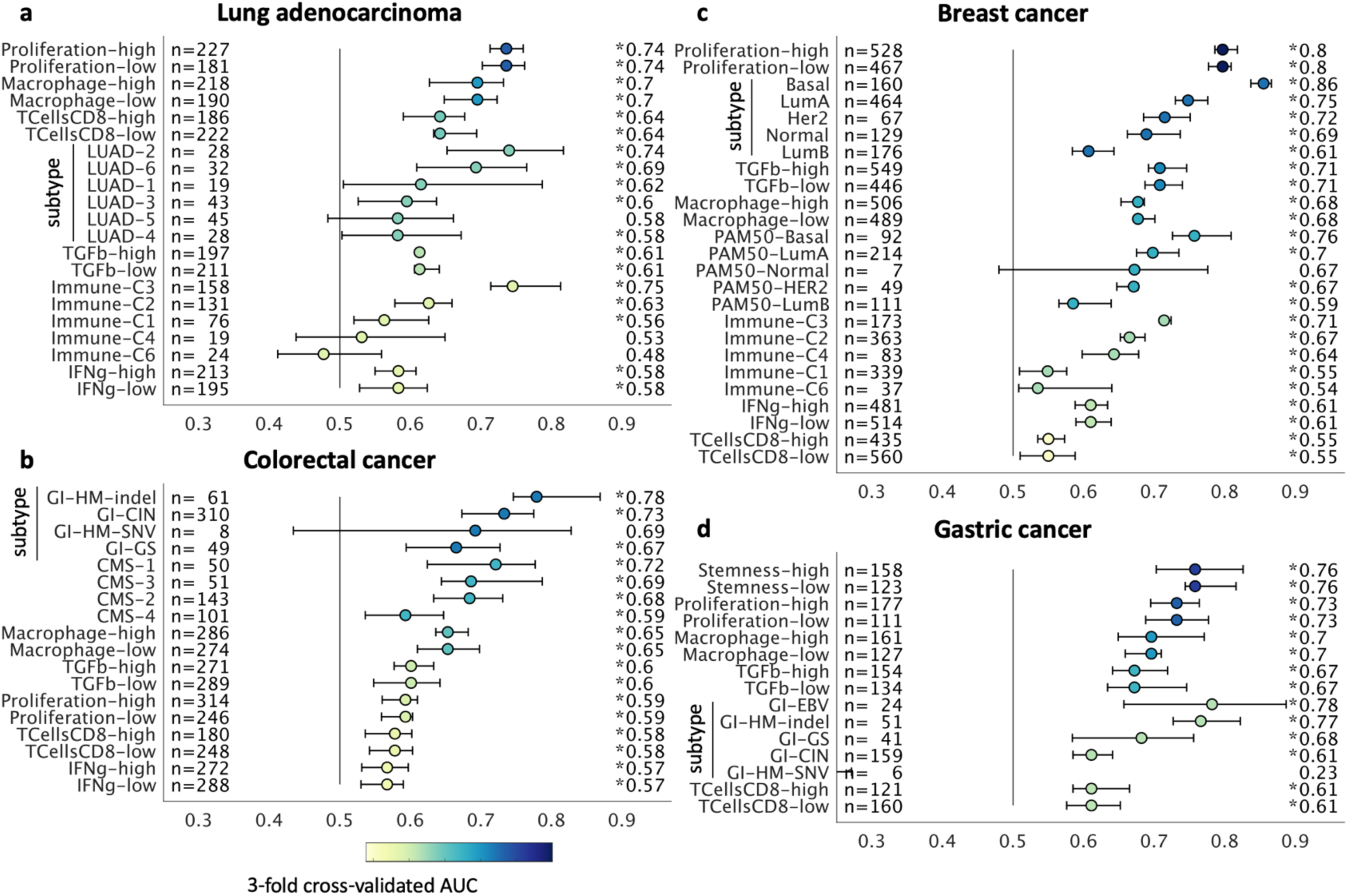
Prediction of gene expression signatures directly from histology. Deep networks were trained to predict clinically relevant gene expression signatures directly from histological images in (**a**) lung adenocarcinoma, (**b**) colorectal, (**c**) breast cancer and (**d**) gastric cancer. Classifiers were assessed by the cross-validated area under the receiver operating curve with bootstrapped confidence intervals (AUC under ROC, horizontal axis). Continuous signatures were binarized at the mean. Variables with an average AUC<0.55 are not shown. (*) denotes all cases where the lower confidence bound exceeds a random classifier (AUC 0.5). “n” denotes the number of patients. For a full list of all tested alterations, see Suppl. Table 2. “subtype” denotes TCGA molecular subtypes.

Finally, we investigated the use of deep learning on conserved molecular classes of tumors such as recently identified TCGA subtypes^3^, pan-gastrointestinal subtypes^4^ and consensus molecular subtypes of colorectal cancer^6^. Few of these classification systems are currently incorporated in clinical workflows, mainly because of the high cost and logistic effort associated with sequencing technology. In our experiments, TCGA molecular subtypes LUAD1-6 were highly detectable in histology images of lung adenocarcinoma (Fig. 3a) with AUCs of up to 0.74. In colorectal cancer (Fig. 3b) and gastric cancer (Fig. 3d), the pan-gastrointestinal (GI) subtypes GI-hypermutated-indel (GI-HM-indel), GI genome stable (GI-GS), GI-chromosomally instable (GI-CIN), GI-hypermu-tated-single-nucleotide variant predominant (GI-HM-SNV) and GI Epstein-Barr-Virus-positive (GI-EBV) were significantly detectable from histology. Correspondingly, in colorectal cancer, ‘consensus molecular subtypes’^6^ were detectable by deep learning (Fig. 3b). These findings could open up fundamentally new options for clinical trials of cancer: While accumulating evidence shows that molecular clusters of tumors are correlated to biologically and clinical outcome, deep molecular classification of these tumors is usually not available to patients in clinical routine or to patients within clinical trials. Detecting these subtypes merely from histology would immediately allow for these subtypes to be analyzed in clinical trials directly from routine material, potentially helping to identify new biomarkers for treatment response. A full description of the methods is available in the “Extended Methods” section.

Together, our results demonstrate the feasibility of pan-cancer deep learning image-based testing. We show that a unified workflow yields reliably high performance across multiple clinically relevant scenarios. Compared to conventional genetic tests, our methodology enables detailed prediction of the spatial heterogeneity of genotypes which is not possible in molecular bulk testing of tumor tissue. An example of this visualization is shown in (Fig. 4a-g): Based only on a routine histological image of colorectal cancer (Fig. 4a), deep learning classifiers correctly predicted CDC27 mutational status (Fig. 4b-c) and consensus molecular subtype (Fig. 4d-g) with a high probability, while assigning a low probability to competing classes.

**Fig. 4.**
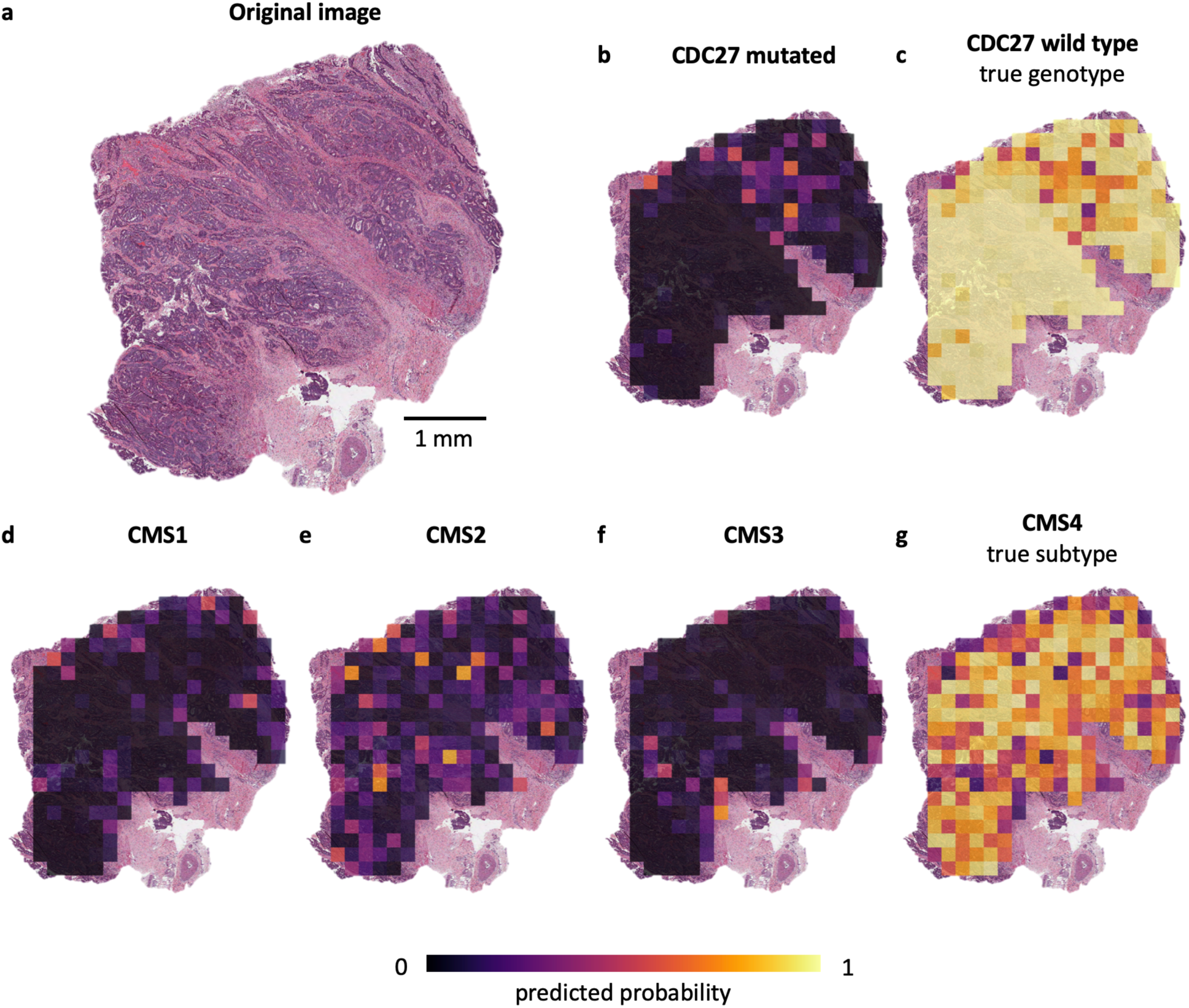
Multiplex genotype maps with local predictability uncovered by deep learning. (**a**) A whole slide image of a colorectal cancer from the TCGA cohort was used for genotype prediction by deep learning classifiers. (**b**) A prediction map for CDC27 wild type status and (**c**) a prediction map for CDC27 mutated status, correctly predicting that this particular patient is mutated. Similarly, prediction maps for consensus molecular subtype (CMS) classes (**d**) CMS1, (**e**) CMS2, (**f**) CMS3 and (**g**) CMS4 correctly show that deep learning robustly predicts CMS from histology alone while highlighting potential intratumor heterogeneity.

Image-based genotyping could be used for definitive testing once performance surpasses previous tests, potentially disrupting clinical workflows Suppl. Fig. 3a-c. A limitation of our method is the low AUC values for some molecular features, but re-training on larger cohorts with up to 10,000 patients per tumor type is expected to increase performance.^16^ Another limitation is that for very unbalanced features – for scarce molecular features – the uncertainty of the AUC estimate is high. Thus, before clinical implementation, multicenter validation is essential, requiring collaborative efforts. Together, our results show that deep learning can consistently unlock dormant patterns in widely available histology images, potentially improving current workflows for molecularly targeted therapy of cancer.

## Author contributions

JNK, ATP and TL designed the study. LH, HIG, NAC, JJS, PAVDB, LFSK and AP oversaw the tumor annotation. CL, AE, JK, HSM, JMN and KAJS manually annotated all tumors. JNK, JK, JMN and PB designed and implemented the algorithm. JNK, CL, AS and NOB curated the list of molecular alterations. HB and MH provided samples from the DACHS study and gave statistical advice. CT, DJ, ATP and TL provided infrastructure and supervised the study. All authors contributed to the data analysis and to writing the manuscript.

## Extended methods

All experiments were conducted in accordance with the Declaration of Helsinki and the International Ethical Guidelines for Biomedical Research Involving Human Subjects. Anonymized scanned whole slide images were retrieved from The Cancer Genome Atlas (TCGA) project through the Genomics Data Commons Portal (https://portal.gdc.cancer.gov/). Tissue samples from the DACHS trial^35,36^ were retrieved from the tissue bank of the National Center for Tumor diseases (NCT, Heidelberg, Germany) as described before.^20^

Scanned whole slide images of tissue slides stained with hematoxylin and eosin were acquired in SVS format. Magnification was between 20x and 40x and corresponding resolution was between 0.25 and 0.51 micrometers per pixel (µm/px). All images were manually reviewed by a trained observer who discussed non-trivial cases with an expert pathologist. After review by the expert pathologist, only those images with tumor tissue on slide were used for downstream analysis. The observer manually delineated tumor tissue on the slide which in most cases included more than half of the total tissue. This region was then tessellated into square tiles of 256 µm edge length. For the benchmark task, these images were resized 1.14 µm/ pixel to be consistent with a previous study^20^; for all subsequent tasks, images were processed at 0.5 µm/pixel. Some patients in the TCGA archive had more than one slide per patient and in these cases, tiles from all slides were pooled on a per-patient basis. From every slide, only a subset of tiles was used for neural network training and prediction (default 1000 tiles per slide; values explored in hyperparameter sampling: 250, 500 and 750). A target variable (e.g. a particular mutation) was matched to each patient (see below) and all tiles corresponding to that patient inherited the label. The patient cohort was then randomly split in three parts in such a way that each part contained approximately the same number of patients with each label. These three parts of the patient cohort were then used for three-fold patient-level cross-validation. Before training, each cohort was randomly undersampled in such a way that the number of tiles per label was identical for each label. For training, we used on-the-fly data augmentation (random x-y-reflection and random horizontal and vertical shear of 5 px). No color normalization was used.

Molecular labels are listed in Suppl. Table 2 and were retrieved from the following sources: Basic clinical and pathological data was retrieved through http://portal.gdc.cancer.gov. Mutational status (wild type or mutated) and high-level amplification were acquired through http://cbiopor-tal.org. In that database, we used “PanCancerAtlas” or “TCGA Provisional” project, whichever contained more patients in that particular tumor type. High-level data on gene expression signatures was retrieved from Thorsson et al. (10). For breast and endometrial cancer, additional data on tumor subtypes were retrieved from Berger et al. (27). For gastric and colorectal cancer, tumor subtype data was retrieved from Liu et al. (11).

Hyperparameter selection was performed for five deep neural networks which were pre-trained on ImageNet: resnet18, alexnet, inceptionv3, densenet201 an shufflenet. The sampled hyperparameter space was as follows: learning rate (fixed) 5e-5 and 1e-4, maximum number of tiles per whole slide image: 250, 500 and 750, number of hot layers (Fig. 1b): 10, 20 and 30. The number of epochs was four with a mini batch size of 512, similar to previous experiments.^20^

All algorithms for whole slide image processing, including tessellation of images and visualization of spatial activation maps, were implemented in QuPath v0.1.2 in Groovy (http://qupath.github.io). All deep learning algorithms, including training and prediction, were implemented in Matlab R2018b (Mathworks, Natick, MA, USA).

All images from the TCGA cohort are available at https://portal.gdc.cancer.gov/. All source codes are available at https://github.com/jnkather/DeepHistology

---

**Suppl. Fig. 1:**
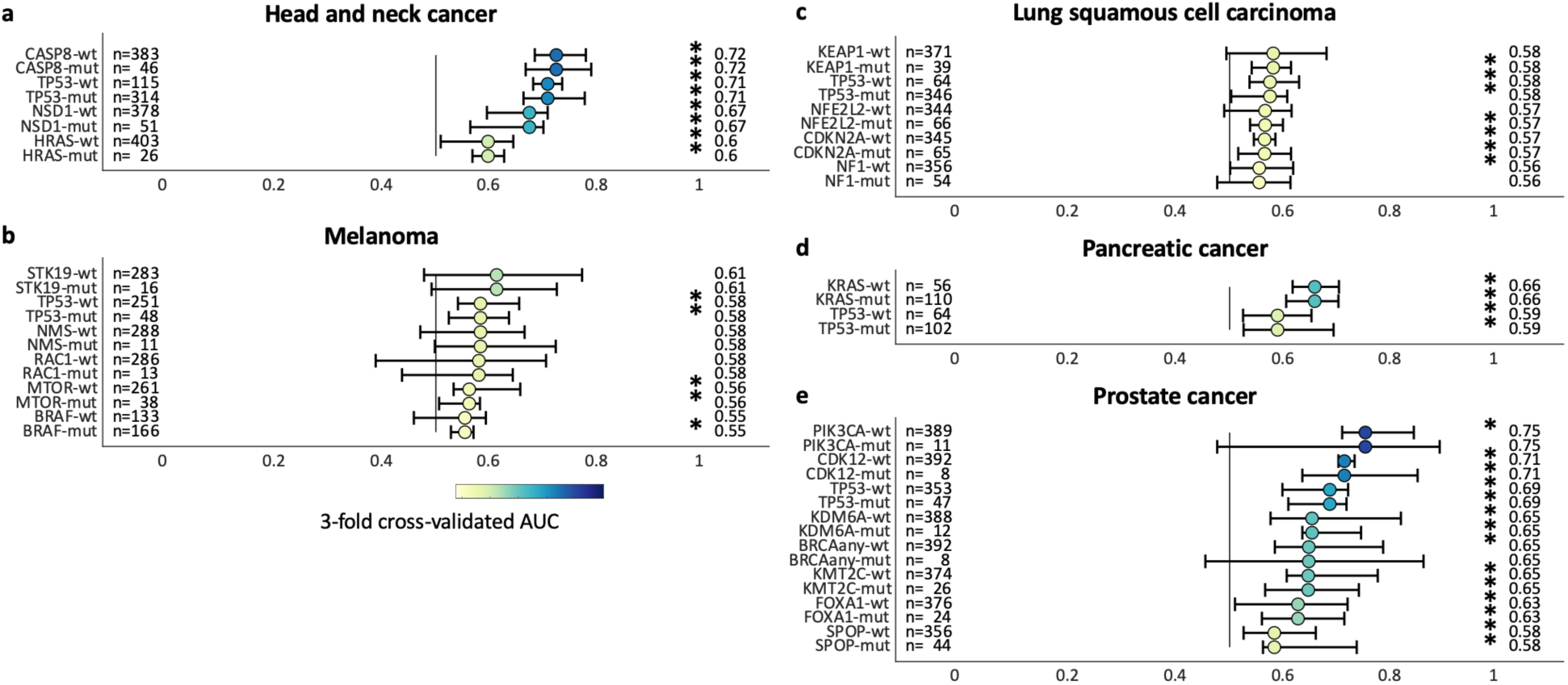
Mutation prediction from histology in additional tumor types. Our method significantly predicted oncogenic mutations from histology in (**a**) Head and neck squamous cell cancer, (**b**) Melanoma, (**c**) Lung squamous cell carcinoma, (**d**) Pancreatic cancer and (**e**) Prostate cancer. The horizontal axis shows three-fold cross-validated area under the receiver operating curve (AUC) as mean +/-95% bootstrapped confidence interval.

**Suppl. Fig. 2:**
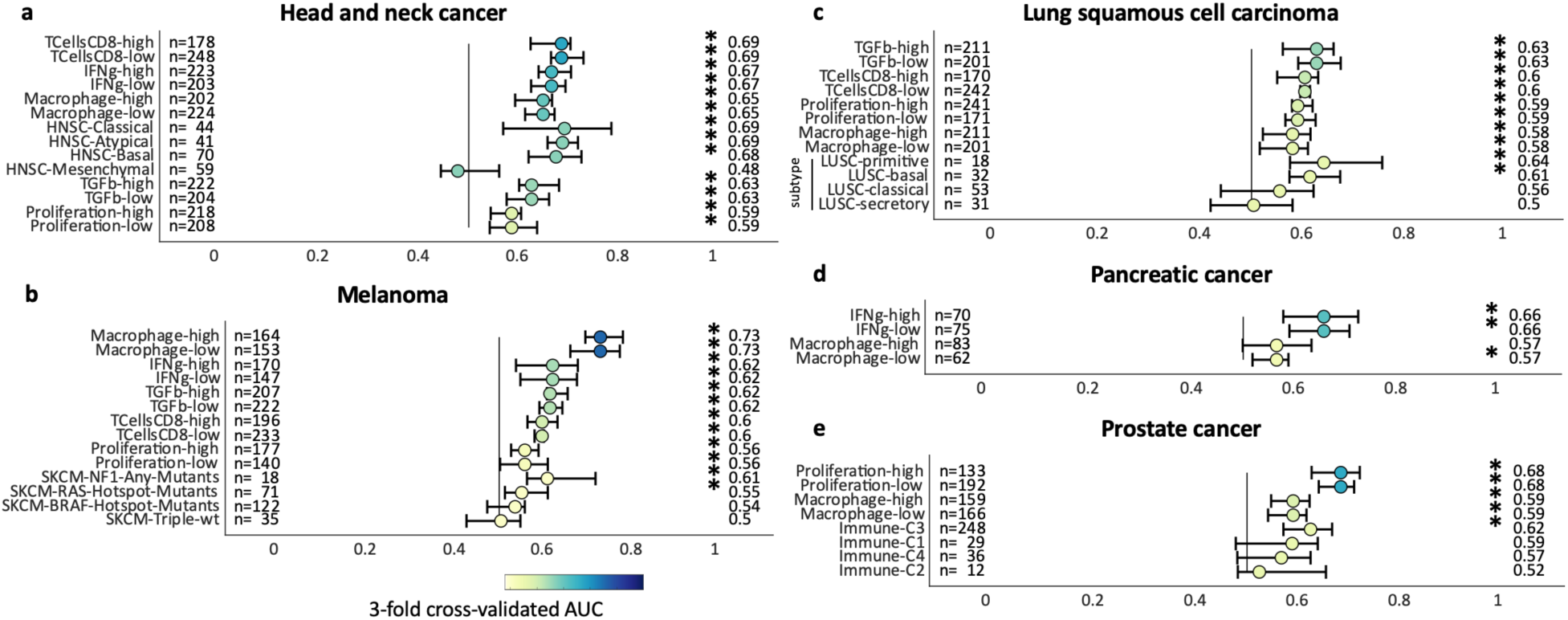
Prediction of high-level gene expression signatures in additional tumor types. Our method significantly predicted high level gene expression signatures from histology in (**a**) Head and neck squamous cell cancer, (**b**) Melanoma, (**c**) Lung squamous cell carcinoma, (**d**) Pancreatic cancer and (**e**) Prostate cancer. The horizontal axis shows three-fold cross-validated area under the receiver operating curve (AUC) as mean +/- 95% bootstrapped confidence interval.

**Suppl. Fig. 3:**
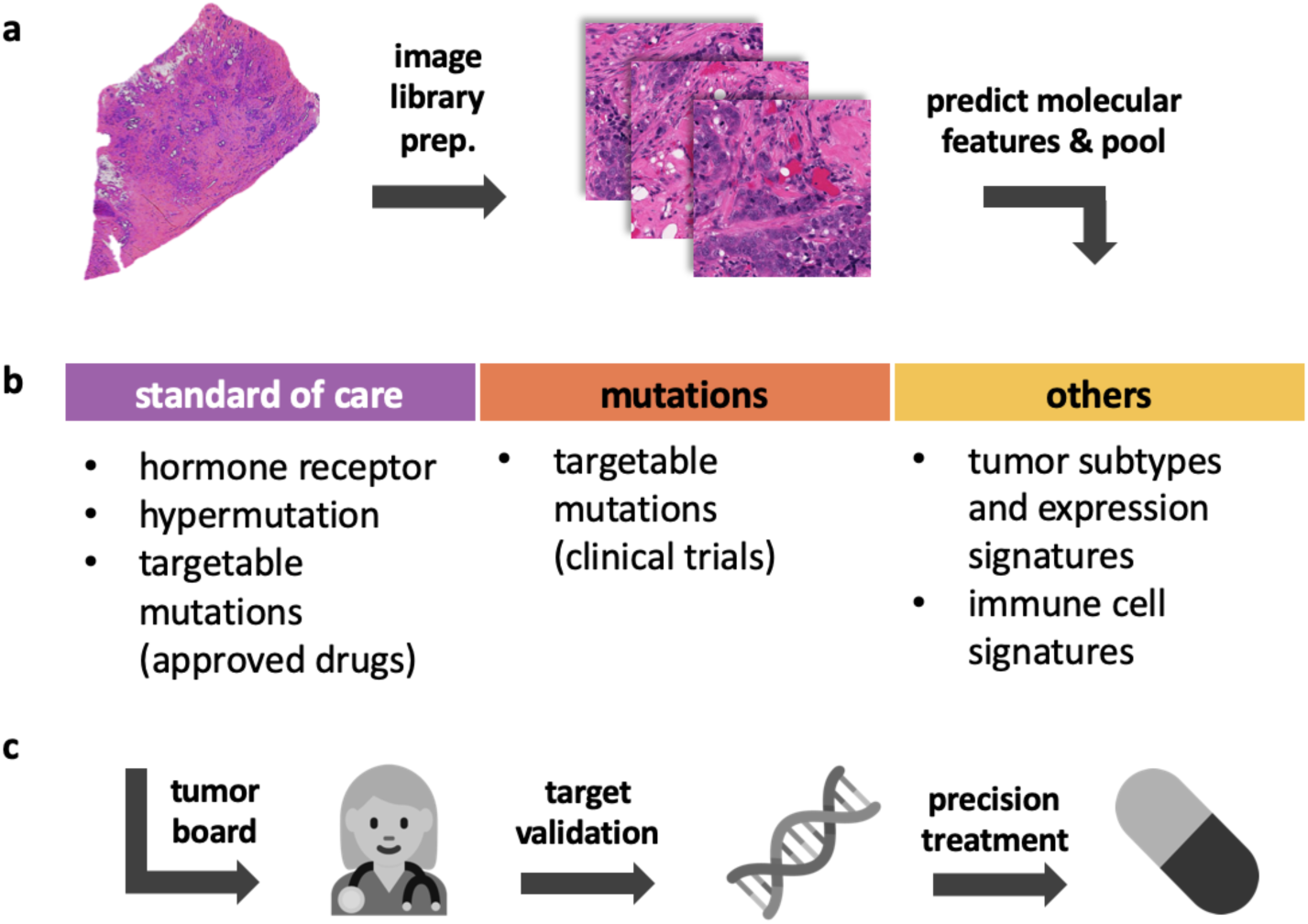
Proposed clinical workflow. (**a**) Starting with ubiquitously available routine histology slides, our method relies on a tessellation of digitized images (“image library preparation”) which are passed to a deep convolutional neural network. The network predicts features on a tile level and the predictions are pooled on a patient level. (**b**) Histology-based testing can be applied to standard of care pathological biomarkers, driver mutations, and other features such as tumor expression subtypes. (**c**) We suggest that clinically meaningful findings of deep learning networks could be discussed in a tumor board, validated by orthogonal methods and ultimately guide targeted treatment.

